# Improving the reliability of fMRI-based predictions of intelligence via semi-blind machine learning

**DOI:** 10.1101/2023.11.03.565485

**Authors:** Gabriele Lohmann, Samuel Heczko, Lucas Mahler, Qi Wang, Julius Steiglechner, Vinod J. Kumar, Michelle Roost, Jürgen Jost, Klaus Scheffler

**Affiliations:** Department of Biomed. Magn. Res. Imaging, University Hospital Tübingen, Germany; Max-Planck-Institute for Biological Cybernetics, Tübingen, Germany; Max-Planck-Institute for Mathematics in the Sciences, Leipzig, Germany; Center for Scalable Dynamics and Artificial Intelligence, Leipzig University, Germany; Santa Fe Institute for the Sciences of Complexity, New Mexico, USA

## Abstract

Predicting neuromarkers for cognitive abilities using fMRI has been a major focus of research in the past few years. However, it has recently been reported that many thousands of participants are required to obtain reproducible results (Marek et al (2022)). This appears to be a major impediment to obtaining neuromarkers from fMRI because large sample sizes are typically not available in neuroimaging studies. Here we show that the out-of-sample prediction accuracy can be dramatically improved by supplementing fMRI with readily available non-imaging information so that reliable predictive modeling becomes feasible even for small sample sizes. Specifically, we introduce a novel machine learning method that predicts intelligence from resting-state fMRI data, leveraging educational level as supplementary information. We refer to our approach as “semi-blind machine learning (SML)” because it operates under the assumption that supplementary information, such as educational level, is available for subjects in both the training and test sets. This setup closely mirrors real-world scenarios, especially in clinical contexts, where patient background information typically exists and can be utilized to boost prediction accuracy. However, guarding against bias is crucial. Subjects should not be categorized as more intelligent simply based on their higher education levels. Therefore, our approach contains a component explicitly designed for bias control. We have applied our method to three different data collections and observed marked improvements in prediction accuracies across a wide range of sample sizes. We anticipate that semi-blind machine learning provides a promising approach to fMRI-based predictive modelling with the potential for a wide range of future applications.

## Introduction

The prediction of individual behaviour from the fMRI-based functional connectome has been investigated in a number of studies, e.g. [1–13], for a review see [14].

However, the viability of this line of research has sparked extensive discussions. Major problems already became apparent at the ADHD-200 Global Competition of 2011 in which researchers were tasked with developing machine learning algorithms for diagnosing ADHD from rsfMRI data [15]. Surprisingly, the team that scored best in this competition did not use any imaging data at all, but instead relied entirely on supplementary non-imaging information [16], see also [17]. And more recently, Marek et al. [18] reported that many thousands of participants are needed to obtain reliable predictions of cognitive abilities using fMRI-based functional connectomes, and even then the prediction accuracies were in fact quite poor. For a debate on this topic see [19–27].

Furthermore, Marek et al. have noted that non-imaging information such as behavioral performance may confound links between brain function and cognitive ability. In fact - much like the ADHD-200 Global Competition - they reported that predictions based on non-imaging information may be more accurate than predictions based on rsfMRI [18, Extended Data Fig 3].

Here we propose to capitalize on such non-imaging information instead of regarding it merely as a confound. Specifically, we introduce a novel machine learning algorithm called “Semi-Blind Machine Learning (SML)” that integrates supplementary non-imaging information into the predictive modelling of the rsfMRI data while carefully controlling potential confounding effects. In essence, our objective is to leverage non-imaging information for improving prediction accuracy while avoiding any potential bias that might be incurred by this process. In SML, the leveraging information is assumed to be known for subjects in both the training and the test set, while the target variable is known only for subjects in the training set. Note that with this design we prioritize prediction accuracy over generalizability. In this work, we will primarily use intelligence as the target variable to be predicted, and educational level as the leveraging factor, but we will also investigate several other target variables.

A straightforward approach of incorporating supplementary information might be to simply average predictions obtained from education levels and neuroimaging data independently. However, as shown in Figure 1, this approach would lead to a strong bias so that subjects with higher education levels would automatically be deemed to be more intelligent. Our goal here is to leverage non-imaging information to improve prediction accuracies while avoiding any such bias. Note that bias in machine learning is a growing concern [28]. In the present context, this issue is particularly relevant because the incorporation of supplementary information would inevitably yield a bias unless countermeasures are taken.

**Figure 1:**
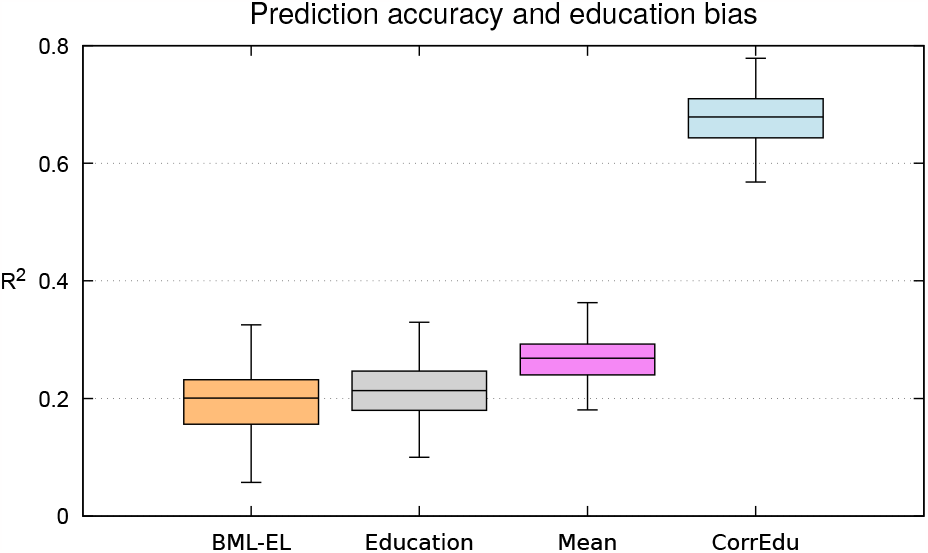
Education bias in straightforward averaging of predictions (HCP). The y-axis shows prediction accuracies for intelligence (“CogTotalComp AgeAdj”) using either just rsfMRI data without additional information about education levels (‘BML-EL’), or just using education levels (‘Education’), or the average of both predictions (‘mean’). The rightmost column shows the variance explained by education of the averaged result. Note that this value greatly exceeds the value shown in the ‘Education’ column. In other words, the averaged prediction is heavily biased so that subjects with higher education levels are automatically deemed to be more intelligent. The rsfMRI data were parcellated using the ICA decomposition into 100 components provided by HCP. Samples sizes for training/test sets were 290 and 100, respectively. 100 random selections of training/test set pairs were used.

Our proposed approach is based on partial least squares regression combined with ensemble learning. The reason for this algorithmic design is that it permits a straightforward means for bias control. We will show a quantitative comparison of our method against ridge regression which is currently one of the most widely used predictive modeling technique in fMRI. Notably, we have excluded neural network approaches from our consideration. This decision is based on recent findings which demonstrate that neural networks do not yield significantly superior performance in this context when compared to conventional regression techniques [29]. Also, our proposed bias control mechanism is not directly transferable to neural network architectures so that a one-to-one comparison would be difficult.

Our method has some similarities with meta-matching [30] in that it also allows predictions with small sample sizes. Meta-matching is based on a transfer learning framework in which a large-scale model is adapted to a separate, smaller sample. A significant improvement in prediction accuracy can be achieved although the large scale model may be trained on a completely different set of phenotypes. This improvement results from inherent correlations among phenotypes, see also [31] for more on phenotypic correlations. In our work, we also harness these phenotypic correlations. However, unlike meta-matching, our enhanced prediction accuracy stems from a novel form of feature selection, rather than from the adaptation of a large-scale model.

### Semi-blind machine learning (SML)

Predictive modelling generally proceeds in three stages [32]. First, brain parcellations are performed to reduce dimensionality. Second, interactions between the brain parcels or components are estimated. Finally, a classifier or regressor is trained to predict behavioural traits or other quantities of interest. The various methods differ with respect to the choice of the strategies used in those three stages, but some form of network modelling is common to all [33, 34]. The methodological challenges and best practices are discussed in [6, 35–40]. In the following, we introduce our Semi-blind Machine Learning method (SML) for predicting intelligence from rsfMRI data.

#### Selecting training and test sets

The first step is to select training and test sets. In our experiments, we randomly selected 100 non-overlapping training/test set pairs of various sizes from a given data collection.

#### Brain parcellations and the connectome

To reduce dimensionality, the fMRI data are parcellated into a set of *k* regions or spatial components. In the experiments reported below, we used several different parcellation schemes including data driven as well as atlas-based methods. For each subject, a connectivity matrix (“connectome”) is computed. Each element of the connectome represents the Pearson linear correlations between fMRI time courses averaged within the parcels or components. Because connectivity matrices are symmetric, we only need to consider the lower triangular parts containing *n* = (*k ×* (*k −* 1))*/*2 edges. As proposed by Varoquaux et al. [41], the parcellated connectomes are embedded into the tangent space. In the following, the term ‘connectome feature’ will be used to denote the edge weight that is computed as the tangent correlation between connectome nodes.

#### Semi-blind ensemble learning (SML-EL)

We have previously proposed a multivariate ensemble learning method to train a prediction model [42]. Here we present an updated version of this method that leverages supplementary non-imaging information – for ease of notation denoted as “L-info”. In the following, we assume that the L-info is available for subjects in both the training and the test set.

We use an ensemble learning approach in which multiple learners are trained independently using different subsets of the feature space [43, 44]. More precisely, a large number of subsets (ensembles) of connectome features are randomly selected, and are then used in a regression model to predict intelligence based on information contained in the training set. This model is subsequently applied to the test set yielding a prediction of intelligence for each subject in the test set. Finally, the predictions from all ensembles are averaged. Here we use partial least squares regression (PLS) because it is particularly well suited to handle problems where the predictors are highly collinear and where the number of independent variables greatly exceeds the number of data points [45–47].

#### Feature selection

In our previous work [42], the connectome features that were recruited for an ensemble were selected randomly. Here we propose to use the L-info to guide the selection so that features that are more likely to be predictive of intelligence are more likely to be included in an ensemble. We call this approach “Semi-blind ensemble learning (SML-EL)”. The term “Blind ensemble learning (BML-EL)” will be used to denote the case when no L-info is used.

In SML-EL, we conduct a univariate analysis on each connectome feature to determine its relevance for predicting IQ. Specifically, for each connectome feature *i* = 1, …, *n*, we calculate the linear correlation *ρ*_*i*_(*train, iq*) between the IQ scores and the corresponding connectome features in the training set. Since IQ scores are unknown in the test set, education is used as a surrogate, and *ρ*_*i*_(*test, edu*) denotes the linear correlation between the education levels and the corresponding connectome features in the test set.

The product of these correlations indicates which connectome features are likely to generalize from the training to the test set:

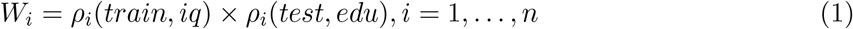

Note that *W*_*i*_ is positive if *ρ*_*i*_(*train, iq*) and *ρ*_*i*_(*test, edu*) have equal signs, and large positive values of *W*_*i*_ indicate a good correspondence between training and test set, provided education is a useful surrogate for intelligence.

In cases where the training set is much larger than the test set, some samples from the training set are randomly selected and transferred to a size-adjusted test set so that the training set and the adjusted test set become equal in size. Note that this is only done for the purpose of feature selection, and a “leakage” from training to test set does not occur. The subsequent feature selection procedure uses the size-adjusted test set.

The weights *W*_*i*_ are now used for a probabilistic feature selection procedure such that features with large weights have a higher probability of being selected into an ensemble. For this, the weights *W*_*i*_ need to be transformed into non-negative weights *P*_*i*_. This transformation is a two-step procedure: first, normalizing the weights through z-scoring, followed by applying a sigmoid function to the normalized values. The sigmoid is defined as

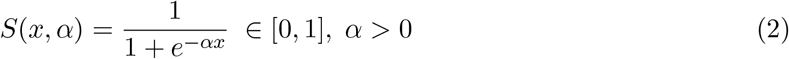

We then use the algorithm by Walker [48] as implemented in [49] for selecting features. More precisely, given features *i* = 1, …, *n* with weights *P*_*i*_ = *S*(*W*_*i*_, *α*), this algorithm produces random indices that are consistent with the given weights so that features with high values of *P*_*i*_ are more likely to be selected.

#### Bias control

The parameter *α* of the sigmoid function is used as a handle for bias control (Figure 2). More precisely, *α* is estimated such that education has a similar correlation with the observed as with the predicted intelligence.

**Figure 2:**
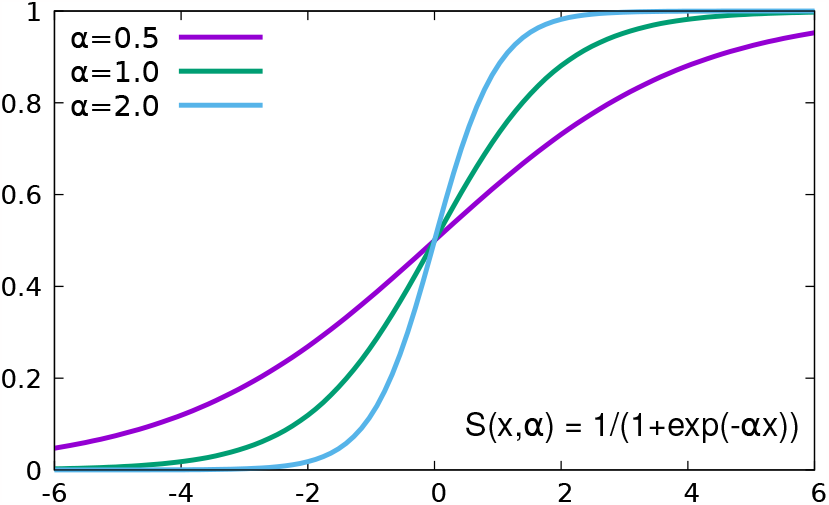
Illustration of bias control. The parameter *α* of the sigmoid function controls the level of randomness in feature selection. Small values of *α* cause features with negative weights to be more likely to be selected into an ensemble so that the level of randomness in the feature selection process is increased, and the impact of education on the final result is reduced.

For this, we compute an initial prediction of intelligence using SML-EL with *α* = 1 as a starting value. We then compute the correlation of education with intelligence scores predicted by SML-EL for the test set, and compare it to the correlation of education with intelligence scores that were observed in the training set. If the correlation in the test set is larger than the correlation in the training set, the parameter *α* is reduced by multiplying it by 0.9, otherwise it is increased by multiplying it by 1.1. This procedure is repeated until convergence, i.e. until the two correlations are sufficiently close. In the experiments reported below, iterations stopped when the two correlations did not differ by more than 0.02.

#### Semi-blind ridge regression (SML-RR)

In addition to ensemble learning, we also implemented ridge regression because it is widely used for predictive modelling and it was shown to have a comparable performance to neural network methods in this and similar tasks [11,29,32]. We used the Scikit-learn package [50, 51] for this purpose. Similar to [52], the regularisation parameter was determined using random search using a three-fold nested-loop crossvalidation.

As in SML-EL, we first conduct a univariate analysis on each connectome feature to determine its relevance for predicting IQ using equation 1. But unlike ensemble learning where a probabilistic feature selection approach was used, we implemented a hard threshold for feature selection. Specifically, the connectome features were ranked using the weights of equation 1, and the top fifty percent were used in the subsequent ridge regression.

## Experimental results

We applied the proposed methods to three different data collections: HCP [53], AOMIC-ID1000 [54] and ABIDE-1 [55]. The HCP data were used for developing the SML-EL algorithm and for finetuning hyperparameters. Note that this may have lead to an algorithmic overfit in the results obtained using HCP.

SML-EL has three hyperparameters, namely the number of ensembles, the size of the ensembles, and the number of PLS components. Using HCP, we found that 1000 ensembles sufficed to reach convergence. The ensemble sizes were set to 1000, and the number of PLS-components were set to 10. We used these three hyperparameter settings in all experiments reported below.

The accuracy of the predictions is reported as the coefficient of determination between the observed and the predicted intelligence defined as

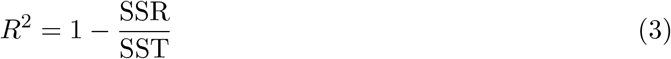

where *SSR* represents the sum of squared residuals and *SST* represents the total sum of squares. In the supplementary material, we also report the results using Pearson linear correlations.

In addition, to estimate potential bias, the variance explained by education is reported. It is computed by fitting a linear regression model using education as the independent variable and the predicted intelligence scores as the dependent variable. The variance explained by education is then defined as in equation 3 with *SST* the variance of the predicted intelligence scores and *SSR* the variance of education levels.

### Human Connectome Project (HCP) [53]

We downloaded minimally preprocessed rsfMRI data of 390 unrelated subjects (202 female, 188 male, aged 22-36, median age 28) acquired at 3 Tesla by the Human Connectome Project (HCP), WU-Minn Consortium (https://db.humanconnectome.org). For dimensionality reduction we used an ICA parcellation into 100 components and the corresponding connectivity matrices containing partial correlations between the components provided by HCP (“netmats2.txt”). The target variable for predictive modelling was the intelligence score provided by HCP called “CogTotalComp AgeAdj”. As L-info we used information about education levels called “SSAGA Educ”. See the supplementary material for more details on these scores and on the scanning and preprocessing parameters of the fMRI data. The results of SML-EL applied to HCP are shown in Figure 3, see also Supplementary Figures S2,S3.

**Figure 3:**
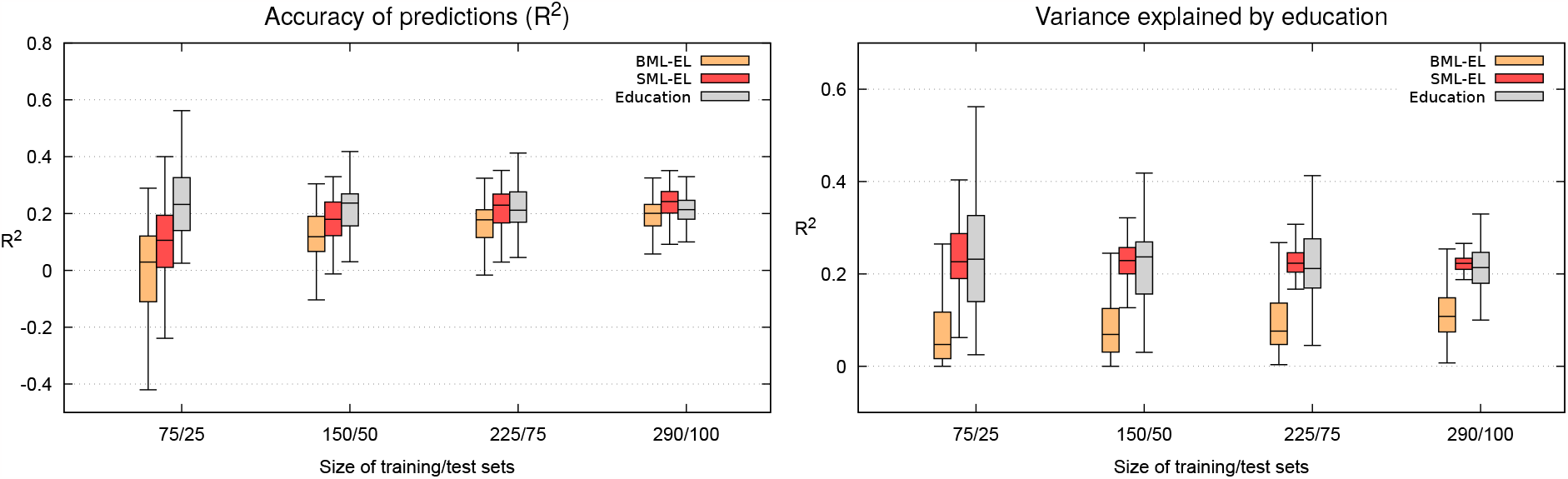
Prediction accuracies using various sample sizes (HCP). Samples sizes for training/test sets are shown along the x-axis. A precomputed ICA-based parcellation using 100 components provided by HCP was used (“netmats2.txt”). The left plot shows prediction accuracies, the right plot shows the variance explained by education. If this value is very different from the variance explained in the observed intelligence, it indicates the presence of bias. Note that the bias control mechanism in SML-EL has prevented such a bias. Results using either blind (‘BML-EL’) or semi-blind ensemble learning (‘SML-EL’) are shown in orange and red. The predictions using education level alone (‘Education’) are shown in gray.

### The Amsterdam Open MRI Collection (AOMIC-ID1000) [54]

We downloaded minimally preprocessed fMRI data of the AOMIC-ID1000 data collection consisting of 928 healthy subjects (https://openneuro.org/datasets/ds003097). For 47 of those, fMRI data were not available on the AOMIC website so that only 881 could be included in our study. Subjects were aged between 19 and 26 yrs (mean 22.8) with 457 females and 424 males.

The target variable for predictive modelling was the intelligence score (‘iq’) as provided by AOMIC. It is based on the Intelligence Structure Test (IST) which measures crystallized and fluid intelligence, as well as memory [57]. We also investigated memory, fluid intelligence, and Body Mass Index (BMI) as further target variables. As auxiliary information we used information about the highest completed educational level (‘edu’) as provided by AOMIC. Here only two different values were recorded corresponding to whether or not participants had completed college. Of the 881 participants, 411 had completed college, all others had not, see [54] for further details.

Figure 4 shows a comparison of semi-blind ensemble learning and ridge regression results using various brain parcellations. The following parcellations were used [58]: dl100(dictionary learning) [56], ica100, [41], 300roi [59], aicha384 [60], mmp360 [61], scha200 [62]. The number of parcels in each parcellation is indicated by their names. Supplementary Figure S4 shows the corresponding results using Pearson linear correlations between observed and predicted intelligence.

**Figure 4:**
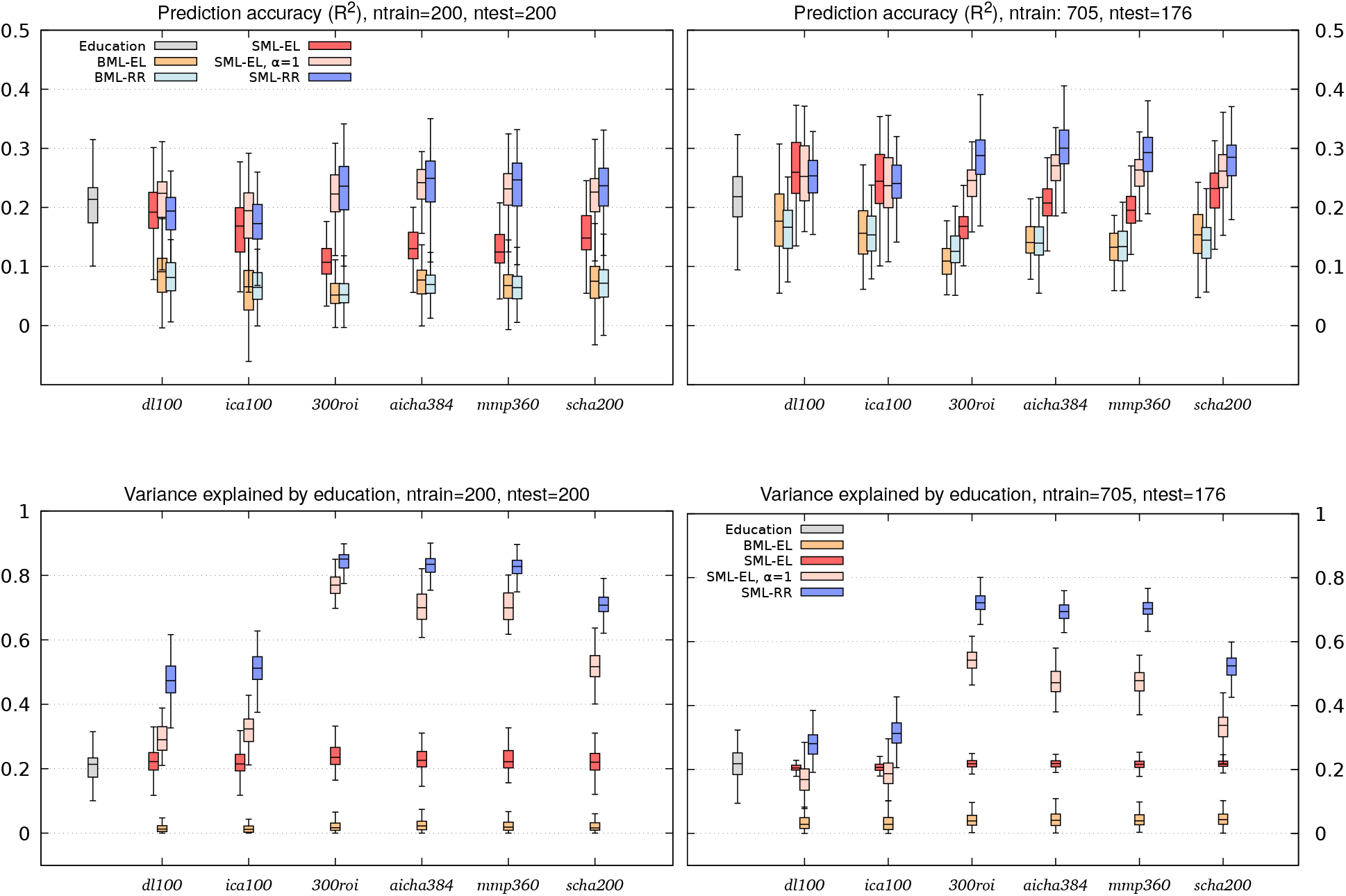
Comparing SML-EL vs SML-RR using various parcellations (AOMIC). The sizes for the training/test sets were 200/200 (left) and 705/176 (right), and 100 random selections of training/test set pairs were used. The top plot shows prediction accuracies, the bottom plot shows the variance explained by education. The number of parcels in each parcellation scheme is indicated by its label, for more details see the main text. Both semi-blind methods SML-RR and SML-EL show a marked improvement in prediction accuracy compared to the blind methods BML-EL and BML-RR. Note that the prediction accuracy is higher in SML-RR compared to SML-EL for some parcellation schemes. However, this comes at the price of a very strong education bias, which is not present in SML-EL. For comparison, the results of SML-EL are also shown without bias control (*α* = 1). Note that in this case, the prediction accuracy is approximately the same as for SML-RR, but with considerably less education bias.

Figure 5 shows results with various sample sizes, see also Supplementary Figure S5. Here, dictionary learning was used for parcellations. Note that even small sample sizes show a clear improvement of prediction accuracy compared to the blind method. Again, SML-RR shows a strong bias, which is not present in SML-EL.

**Figure 5:**
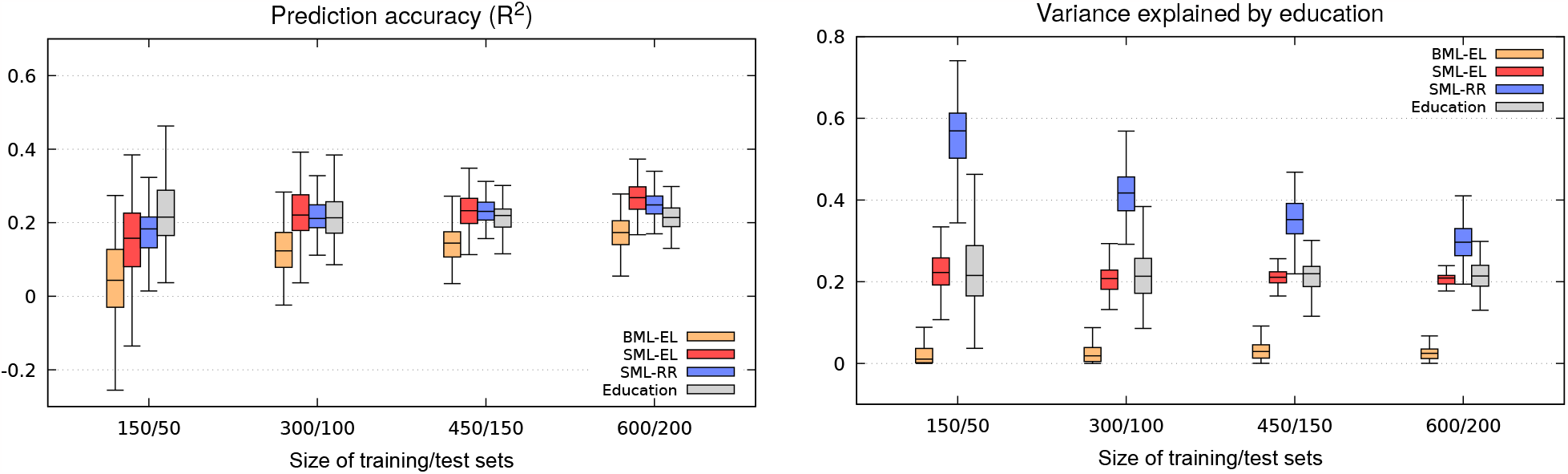
Accuracy of prediction using various sample sizes (AOMIC). Samples sizes for training/test sets are shown along the x-axis. 100 random selections of training/test set pairs were used. The data were parcellated into 100 parcels using dictionary learning [56]. The left plot shows prediction accuracies, the right plot shows the variance explained by education. Note that SML-RR is strongly biased by education, whereas SML-EL does not show a bias. Also note that BML-EL shows a clear negative bias, i.e. the correlation of education with the predictions of IQ in BML-EL is much lower than the correlation of education with the observed IQ.

Figure 6 was included to show the effect of increasing training set sizes. Note that the prediction accuracy initially correlates with increases in sample size but reaches a plateau at about 600, indicating that larger sample sizes may not be sufficient to improve prediction accuracies, see also Supplementary Figures S6,S7.

**Figure 6:**
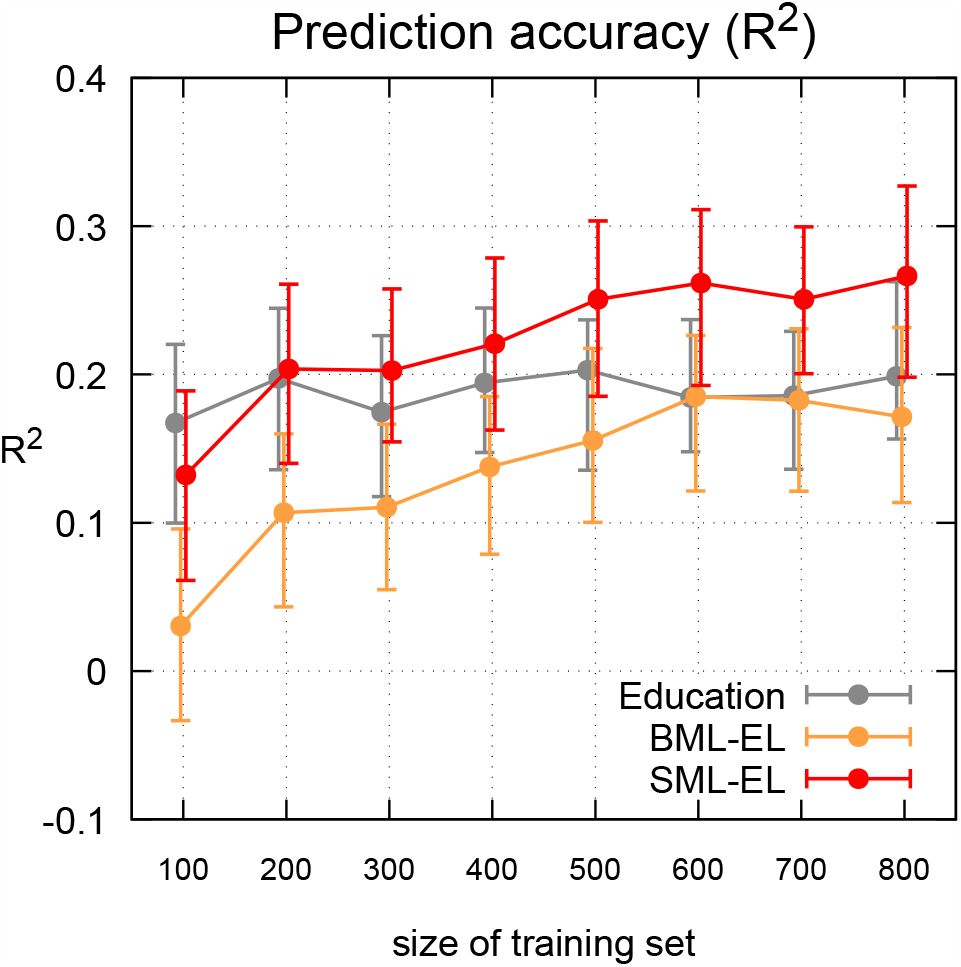
Prediction accuracy reaches a plateau (AOMIC). The size of the training sets are shown on the x-axis. The size of the test sets was 80 throughout. The error bars show the upper and lower quartiles. Note that the prediction accuracy initially correlates with the size of the training sets, but reaches a plateau at a sample size of about 600 beyond which it remains relatively constant. Also note that the semi-blind approach (SML-EL) surpasses the prediction accuracy of “Education” for training sample sizes larger than 300, whereas the blind approach (BML-EL) fails in this regard. As before, the data were parcellated into 100 parcels using dictionary learning [56], see also Supplementary Figures S6,S7.

In order to further investigate the role of the L-info for prediction accuracy, we also used simulated L-info and applied it to four different target variables, namely IQ, fluid intelligence, memory and Body Mass Index (BMI) as provided by AOMIC. The simulated L-info was randomly generated with several pre-defined levels of correlation with the target variable ranging from 0.3 to 0.6. The results are shown in Figure 7 and Supplementary Figures S8,S9. Note that even a weak correlation of the simulated L-info with the target variable helps to improve prediction accuracy.

**Figure 7:**
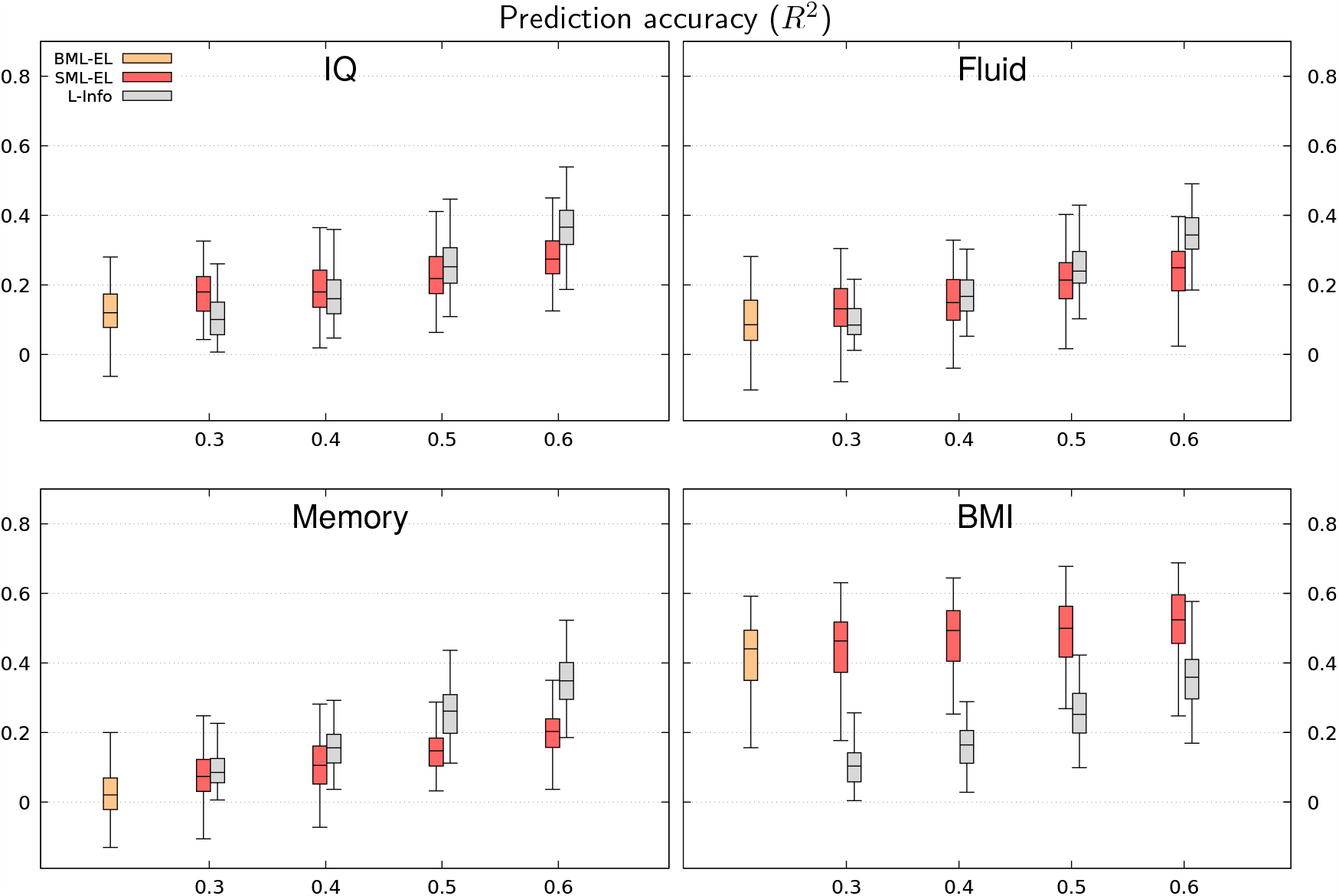
Predictions of various target variables using simulated leverage information (*R*^2^) - AOMIC. This figure shows predictions of IQ, fluid intelligence, memory and body mass index (BMI) of the AOMIC collection. The leverage information was randomly generated to various degrees of linear correlation with the target variable as indicated by the numbers on the x-axis (‘0.3’,’0.4’,’0.5’,’0.6’). The sizes of training/test sets were 300/100. Note that even weakly correlated L-info helps to improve prediction accuracy. Using strongly correlated L-info alone (without fMRI) may yield higher prediction accuracies than SML-EL, but the results would of course be heavily biased.

### Autism Brain Imaging Data (ABIDE-1) [63]

The ABIDE-1 data collection is typically used to investigate autism spectrum disorders (https://fcon_1000.projects.nitrc.org/indi/abide/abide_I.html). For the present study we only included data of 442 healthy controls aged 6 to 56 yrs (mean 16.4), while patient data were excluded, see the supplementary material for more details. The reason why we included this specific collection is its greater heterogeneity in comparison to the other two collections. This increased heterogeneity is due to a much wider age range of participants and the data being collected from 16 distinct sites. In this scenario, model prediction becomes significantly more challenging, see also [11]. The target variable for predictive modeling was the FIQ Standard Score (full IQ) as provided by ABIDE-1, and also subject age. Unfortunately, ABIDE-1 does not contain information about education levels. Therefore, we randomly generated data to simulate supplementary non-imaging information with a pre-determined correlation with FIQ and age. Brain parcellations were done using the Dosenbach-160 Atlas [64]. The results of SML-EL applied to ABIDE-1 is shown in Figure 8, see also Supplementary Figures S10,S11. As for the other two data collections, the L-info provided increases in prediction accuracy compared to the blind versions of the algorithm. This was true for both target variables, even though the prediction accuracy for age is markedly better than for FIQ.

**Figure 8:**
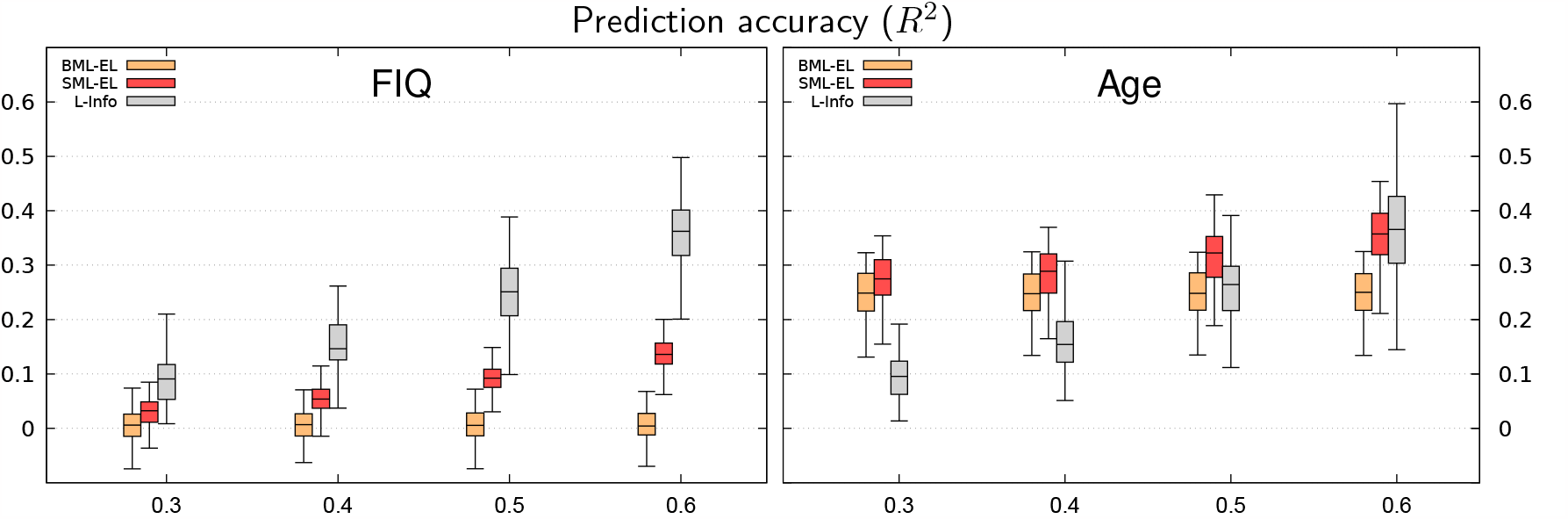
Predictions of various target variables using simulated leverage information (*R*^2^) - ABIDE-1. This figure shows predictions of FIQ and age of the ABIDE-1 collection. The leverage information was randomly generated to various degrees of linear correlation with the target variable as indicated by the numbers on the x-axis (‘0.3’,’0.4’,’0.5’,’0.6’). The sizes of training/test sets were 300/100.

## Discussion

We have introduced a semi-blind machine learning approach designed to predict intelligence from fMRI data. By incorporating education as leveraging information, we achieved a significant improvement in out-of-sample prediction accuracy. Moreover, we found that sample sizes as small as a few hundred were sufficient to attain prediction accuracies that would typically require thousands of participants. This implies a promising avenue for addressing the concerns raised by Marek et al [18].

As shown in Figure 6, relying solely on increasing sample sizes may not be sufficient for resolving the issue of inadequate prediction accuracy. Therefore, we argue that completely new approaches like the one presented in this paper are needed. Here we have proposed to leverage non-imaging supplementary information for boosting prediction accuracy. We have introduced a bias control mechanism that guarantees that the resulting predictions maintain a correlation with the non-imaging information similar to that observed in the target variable within the training dataset. Without such a mechanism, the resulting predictions would be heavily biased.

We are aware that our semi-blind models are not generalizable to individuals outside of the given test set because they leverage information from the test set. But we argue it is uncertain whether any model can genuinely claim to be universally applicable to the entire human population so that generalisability may not be a viable objective in any case. Furthermore – because computation times are on the order of a few minutes – semi-blind models can be easily updated if a new set of test subjects arrives.

As noted in the introduction, our approach is related to meta-matching [30] in that it also aims for reducing the sample sizes needed for reliable predictions. But it differs in that it does not require a large-scale model trained on thousands of subjects. Another key distinction is our explicit bias control mechanism which is not included in meta-matching.

We have proposed two different semi-blind machine learning methods both showing a marked improvement of prediction accuracy. The advantage of ensemble learning (SML-EL) combined with a probabilistic feature selection is that it offers a straightforward way of controlling the bias. Ridge regression on the other hand (SML-RR) was found to be considerably more vulnerable to bias.

As expected, we found that information about educational levels is highly predictive of intelligence and clearly outperforms conventional fMRI-based regression methods. However, our proposed semi-blind approach was able to outperform predictions made solely based on education, particularly when using parcellations derived from dictionary learning or ICA. Furthermore, it offers the advantage of avoiding bias, resulting in predictions that reflect intelligence rather than merely educational levels. Additionally, it provides neuroimaging data that can be used to identify brain regions associated with cognitive abilities.

We found that the parcellation schemes with fewer parcels (*ica100, dl100*) produced better results than more fine-grained parcellations (*mmp360, scha200, aicha384*). This is likely due to the fact that a semi-blind feature selection is more challenging if more features are available. Another explanation might be that *ica100* and *dl100* are both data-driven and are therefore optimized for the data collection at hand. However, in our experiments we have used small subsets of data while these two parcellations were in fact based on much larger assemblies.

In summary, we conclude that semi-blind machine learning provides a promising approach to fMRI-based predictive modelling that lends itself to a wide range of potential future applications.

## Software

The software will be made public soon.

## Acknowledgments

This work was partially funded by DFG-SPP 2041 “Computational Connectomics”.

Samuel Heczko was supported by the International Max Planck Research School for the Mechanisms of Mental Function and Dysfunction (IMPRS-MMFD).

Data were provided by the Human Connectome Project, WU-Minn Consortium (Principal Investigators: David Van Essen and Kamil Ugurbil; 1U54MH091657) funded by the 16 NIH Institutes and Centers that support the NIH Blueprint for Neuroscience Research; and by the McDonnell Center for Systems Neuroscience at Washington University.

