## Supplementary Material for "Improving the reliability of fMRI-based predictions of intelligence via semi-blind machine learning"

### Human Connectome Project (HCP) [1]

The rsfMRI data were acquired in two sessions on two separate days with two different phase encoding directions (left-right and right-left) with spatial resolution $(2mm)^3$, multiband factor 8. Each scan hat 1200 volumes acquired at TR=0.72 seconds so that the total scan time across all four sessions was approximately 58 minutes. We excluded data sets for which data quality problems due to instability of the head coil were reported [2]. In our study we used data of 390 unrelated subjects with 188 males, 202 females. They were aged between 22 and 36 yrs, mean (28.5 yrs).

We used the network matrices provided by HCP that were derived from ICA decompositions into 100 components with partial correlations between the components ("netmats2.txt"). For further details, see

`https://www.humanconnectome.org/storage/app/media/documentation/s1200/HCP1200-DenseConnectome+PTN+Appendix-July2017.pdf`

As supplementary information, we used information about education levels provided by HCP called ('SSAGA_Educ'). It records the number of years of education completed ranging from 11 to 17 yrs (mean 15 yrs). Below, a histogram of the 390 subjects used in our study is shown.

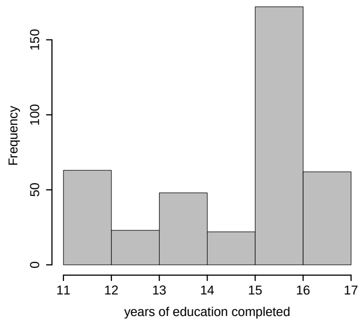


**Supplementary Figure S1: Histogram of supplementary information 'SSAGA_Educ'.**

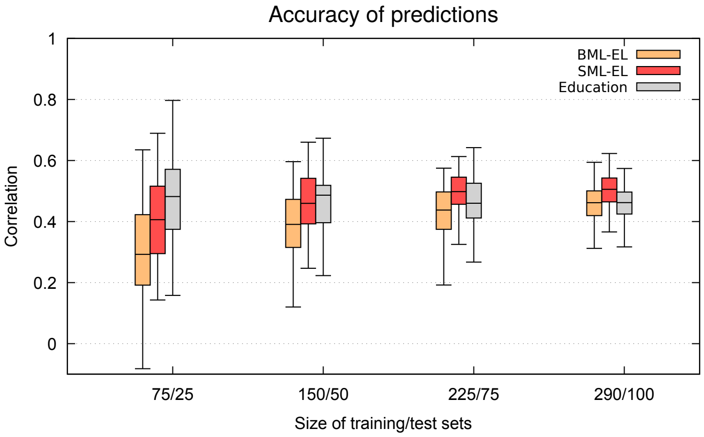


**Supplementary Figure S2: Prediction accuracies using various sample sizes (HCP).** *This figure corresponds to the lefthand plot Figure 3 of the main text. It shows correlations (rather than $R^2$) between observed and predicted intelligence.*

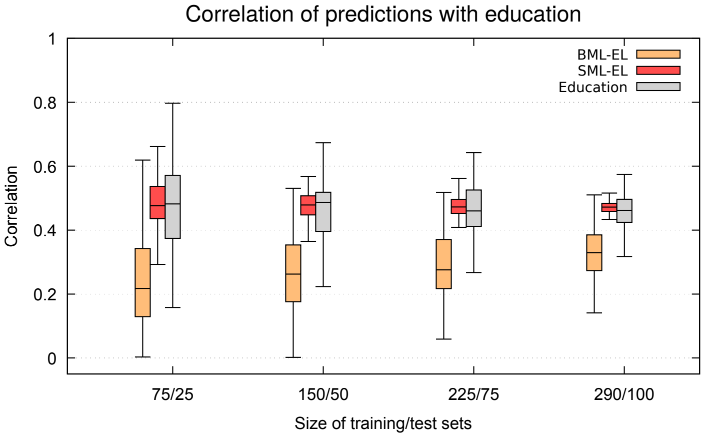


**Supplementary Figure S3: Correlations with education (HCP).** *This figure corresponds to the righthand plot of Figure 3 of the main text. It shows correlations between predicted intelligence and education levels.*

### The Amsterdam Open MRI Collection (AOMIC-ID1000) [3]

Data were acquired on a Philips 3T Intera scanner (Philips, Best, the Netherlands). The duration of the fMRI scan was approximately 11 minutes (290 volumes) with TR=2.2 seconds. The spatial resolution was $3 \times 3 \times 3.3$mm. We used minimally preprocessed data provided on the AOMIC website called: "sub-????_taskmoviewatching_space-MNI152NLin2009cAsym_desc-preproc_bold.nii.gz".

The data were corrected for baseline drifts using a highpass filter with a cutoff frequency of 0.1 Hz and a spatial Gaussian filter with fwhm=6mm was applied. As described in [3], subjects viewed a movie clip during the fMRI scan consisting of a compilation of 22 natural scenes.

We investigated several parcellations as listed below. They are referenced in Figure 4 of the main manuscript.

| Parcellation | ROIs | reference |
|---:|:---:|:---:|
| dl100 | 100 | [4] |
| ica100 | 100 | [5] |
| 300roi | 300 | [6] |
| aicha384 | 384 | [7] |
| mmp360 | 360 | [8] |
| scha200 | 200 | [9] |

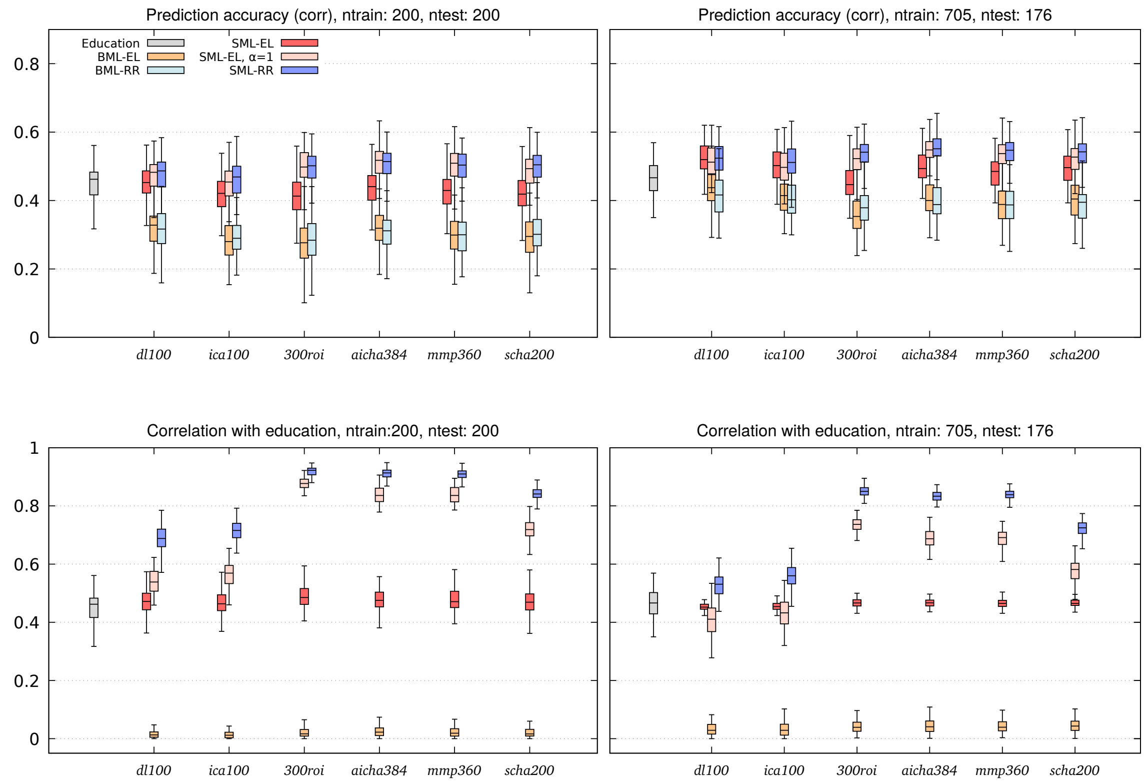


**Supplementary Figure S4: Comparing various parcellations (AOMIC).** *This figure corresponds to Figure 4 of the main text. It shows correlations (rather than $R^2$) between observed and predicted intelligence.*

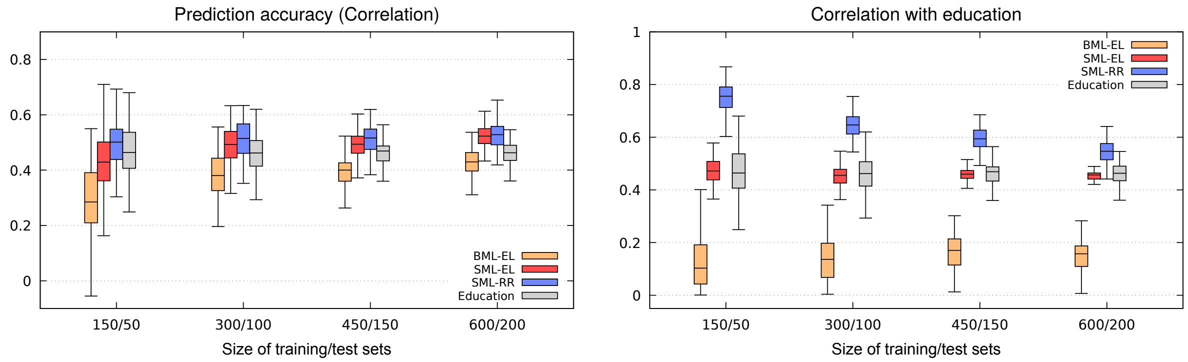


**Supplementary Figure S5: Prediction accuracies using various sample sizes (AOMIC).** *This figure corresponds to Figure 5 of the main text. It shows correlations (rather than $R^2$) between observed and predicted intelligence.*

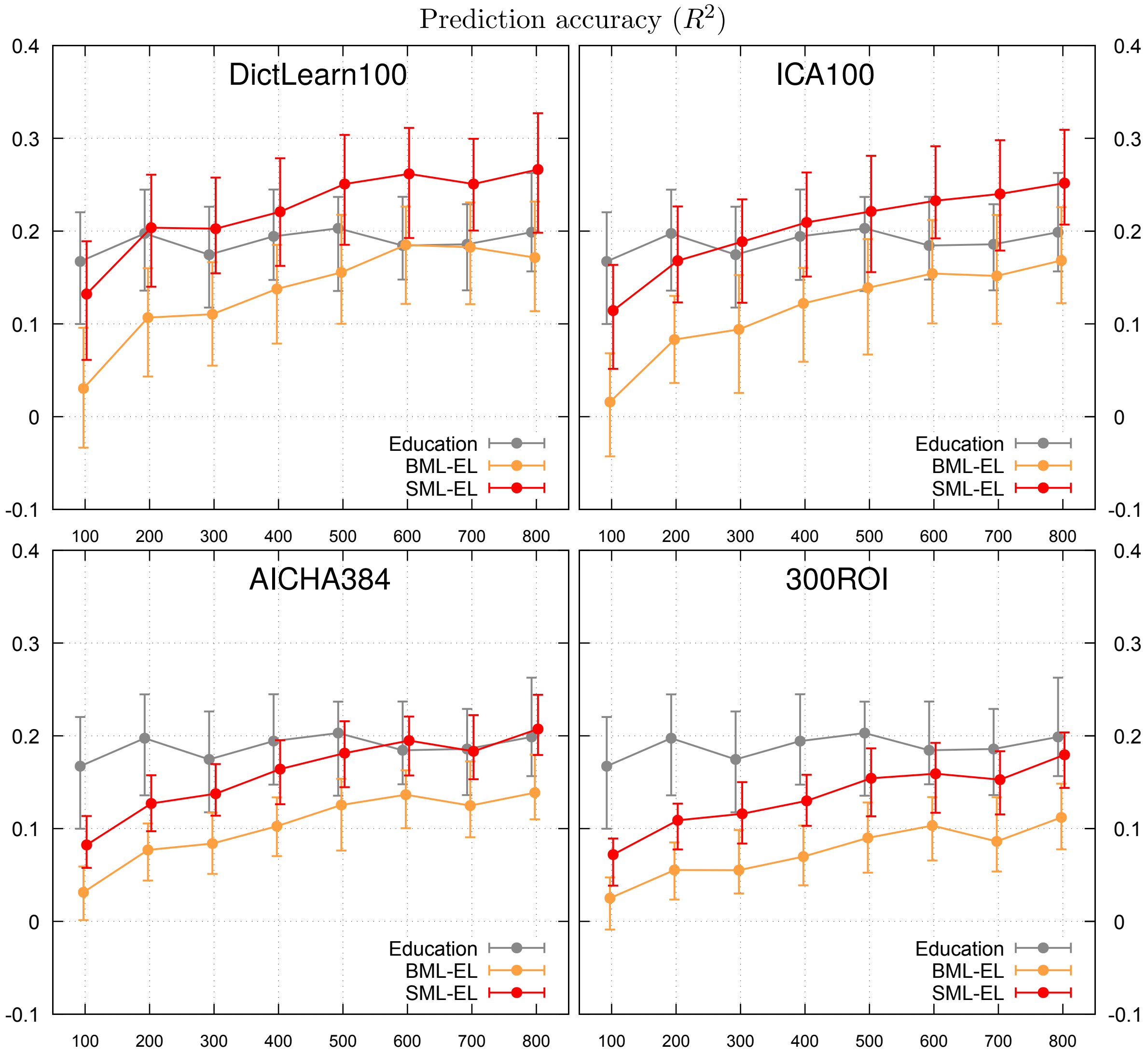


**Supplementary Figure S6: Prediction accuracy across various sizes of the training set (AOMIC).** *The size of the test set was 80 throughout. Prediction accuracies ($R^2$) for four different parcellation scheme are shown. Note that for dictionary learning and ICA, semi-blind regressions surpasses "Education" if training sample sizes exceed about 300, the other two parcellation schemes are less successfull. Also note that for Dictionary Learning and ICA, prediction accuracy seems to reach a plateau at a sample size of about 600.*

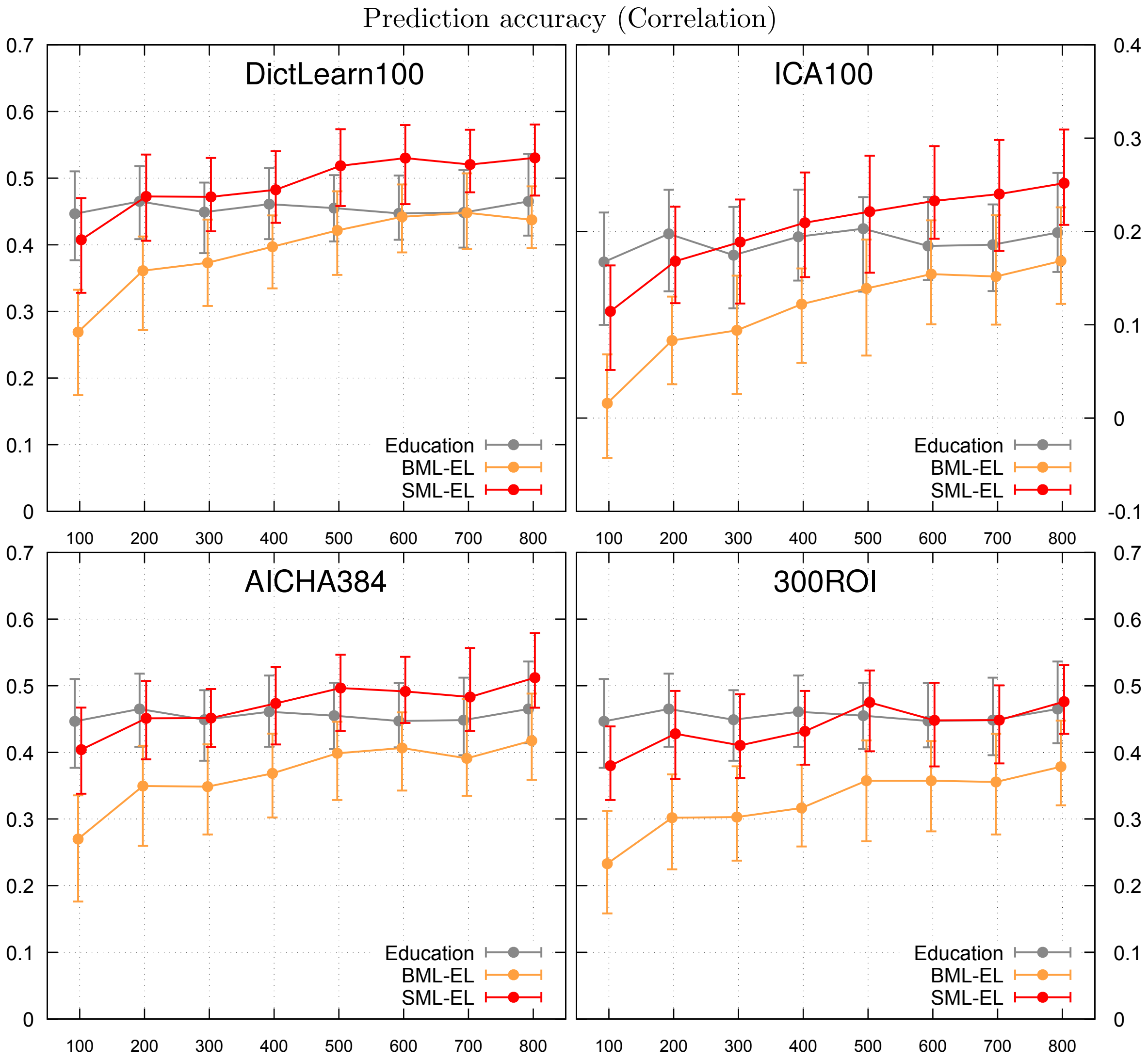


**Supplementary Figure S7: Prediction accuracy across various sizes of the training set (AOMIC).** *Similar to Supplementary Figure S6, but here prediction accuracies are reported as Pearson linear correlations between observed and predicted intelligence.*

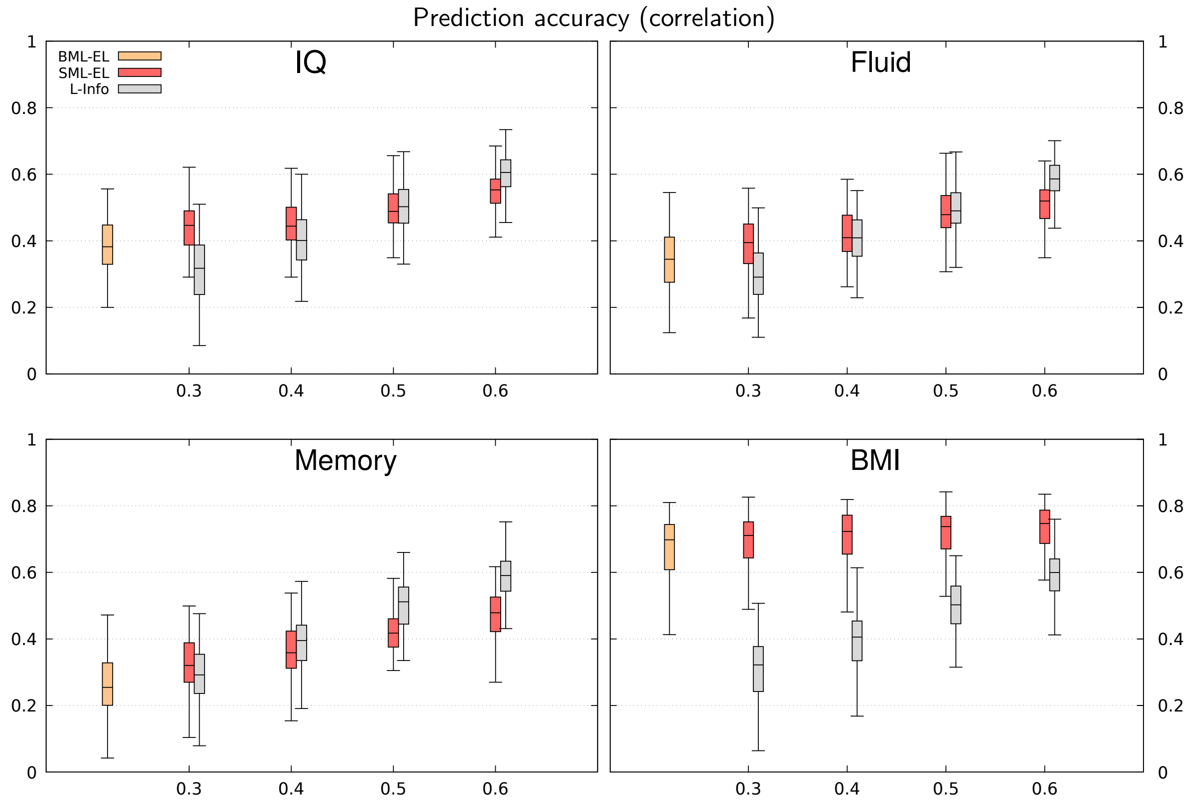


**Supplementary Figure S8: Predictions using simulated L-Info (AOMIC).** *This figure corresponds to Figure 7 of the main text. It shows correlations between observed and predicted target variables. The L-info was generated as described in the main text. The training/set set sizes were 300/100.*

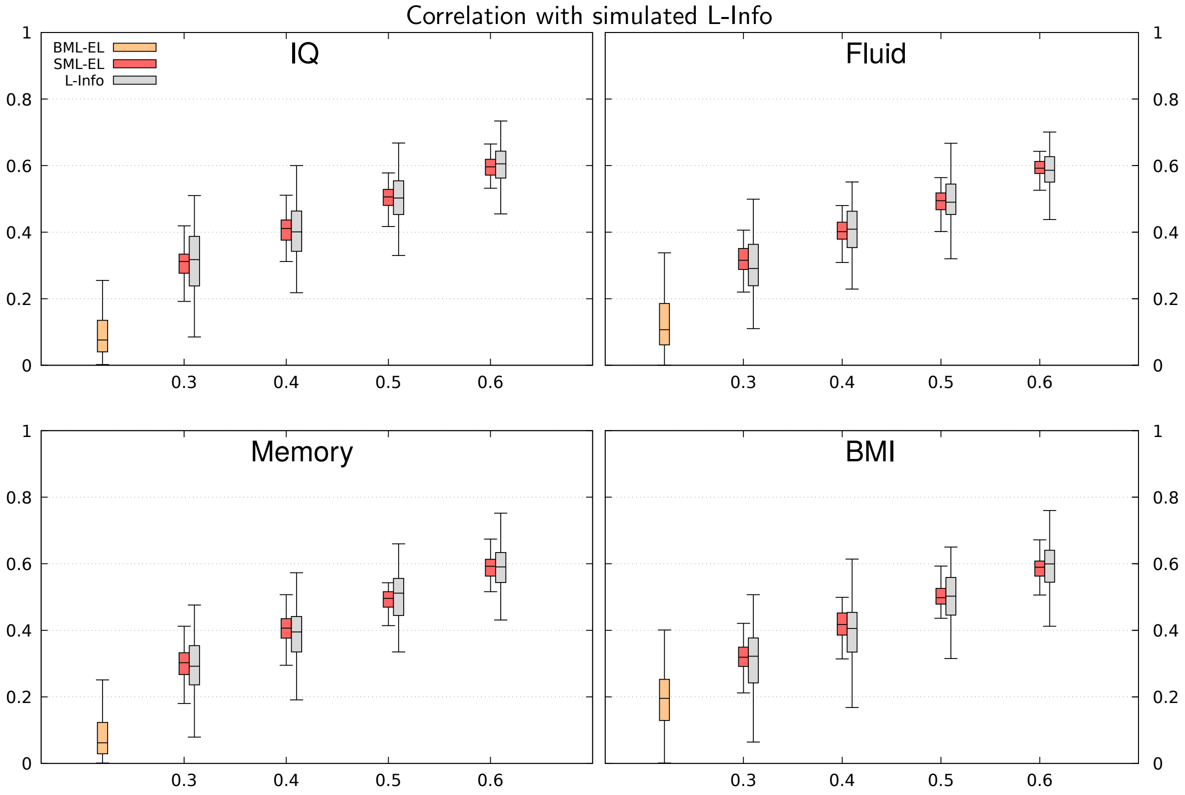


**Supplementary Figure S9: Correlation with simulated L-Info (AOMIC).** *This figure corresponds to Figure 7 of the main text. It shows correlations between predicted targets and L-info.*

### Autism Brain Imaging Data (ABIDE-1)

The data were preprocessed by the Preprocessed Connectomes Project (PCP) [10] using the following steps.

1. SPM8 based DPARSF pipeline
2. discard first 4 volumes of each fMRI time series for magnetization stabilization
3. slice timing correction
4. realignment to first volume to account for head motion
5. no intensity normalization
6. 24-parameter head motion, mean white matter and CSF signals regressed out
7. motion realignment parameters, linear and quadratic trends in low-frequency drifts are regressed out
8. bandpass filtering (0.01 - 0.1Hz) after nuisance signal regression
9. functional to anatomical registration using rigid body transformation
10. anatomical to standard space (MNI152) registration using DARTEL
11. smoothing with 6mm FWHM Gaussian kernel
12. resampling to a spatial resolution of $(3mm)^3$.

**Computing functional connectivity:**

For each subject:

1. parcellations using the Dosenbach-160 Atlas [11].
2. calculate PCC using nilearn.connectome.ConnectivityMeasure with the following parameters: kind="correlation", vectorize=True, discard_diagonal=True

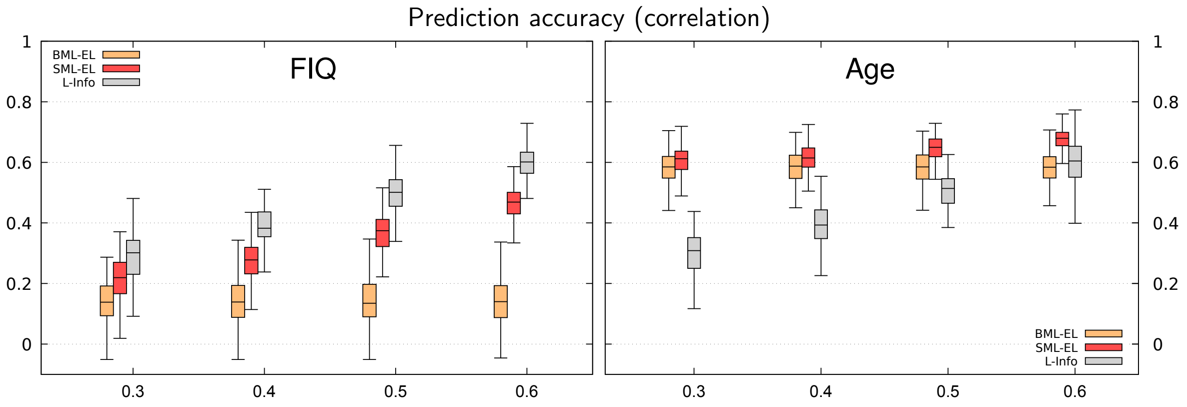


**Supplementary Figure S10: Predictions using simulated L-Info (ABIDE-1).** *This figure corresponds to Figure 8 of the main text. It shows correlations between observed and predicted target variables. The L-info was generated as described in the main text. The training/set set sizes were 300/100.*

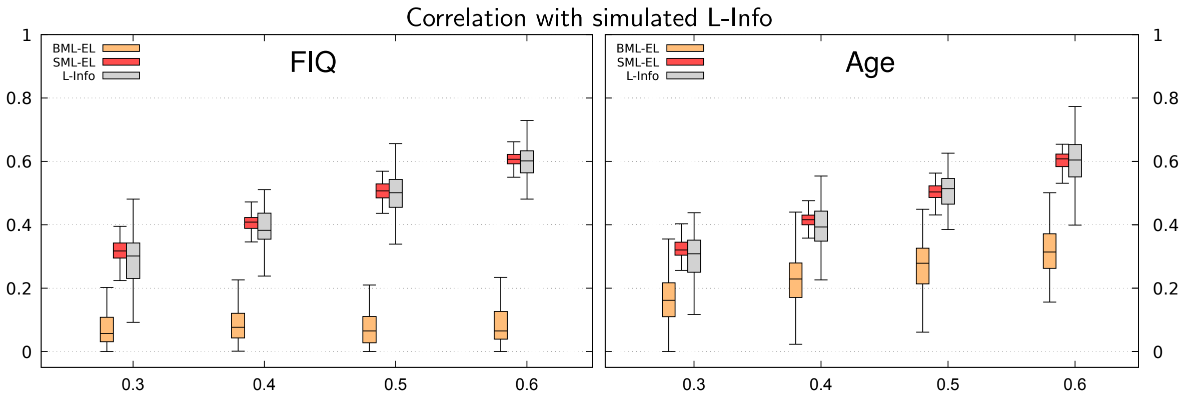


**Supplementary Figure S11: Correlation with simulated L-Info (ABIDE-1).** *This figure corresponds to Figure 8 of the main text.*
